# Methods for cell isolation and analysis of the highly regenerative tunicate *Polycarpa mytiligera*

**DOI:** 10.1101/2023.07.21.549655

**Authors:** Tal gordon, Noam Hendin, Omri Wurtzel

**Affiliations:** The School of Neurobiology, Biochemistry & Biophysics, George S. Wise Faculty of Life Sciences, Tel Aviv University, Tel Aviv, Israel; Sagol School of Neuroscience, Tel Aviv University, Tel Aviv, Israel

## Abstract

*Polycarpa mytiligera* is the only molecularly characterized solitary ascidian capable of regenerating all organs and tissue types. The cellular basis for regeneration in *P. mytiligera* is largely unknown, and methods for isolating live cells from this species for functional analyses are unavailable. Here, we developed a method for isolating live cells from *P. mytiligera*, overcoming major experimental challenges, including the dissociation of its thick body wall and native cellular autofluorescence. We demonstrated the applicability of our approach for tissue dissociation and cell analysis using three flow cytometry platforms, and by using broadly used non-species-specific cell labeling reagents. In addition to live cell isolation, proof-of-concept experiments showed that this approach was compatible with gene expression analysis of RNA extracted from the isolated cells, and with ex vivo analysis of phagocytosis. The ability to purify live cells will promote future studies of cell function in *P. mytiligera* regeneration.

## Introduction

Ascidians (Phylum: Chordata, Class: Ascidiacea) are marine filter feeding chordates, and are part of the sister group of vertebrates^1^. Ascidians are broadly used in ecological^2–4^, cellular^5,6^, and molecular research^7^. Colonial ascidians reproduce both sexually and asexually and can regenerate any missing body part^8,9^. By contrast, solitary ascidians only reproduce sexually and have a varying regeneration capacity^10,11^. *Polycarpa mytiligera* is the only known solitary ascidian species that can regenerate any body tissue, making it an attractive candidate for functional studies of regeneration^12^. Recent studies have characterized regeneration in *P. mytiligera* and have established key techniques for molecular analysis: (1) production of a transcriptome assembly^13,14^; (2) characterization of gene expression during regeneration^13^ and injury response^14^; (3) optimization of methods for analyzing gene expression by fluorescent in situ hybridization (FISH) or immunofluorescence in tissue sections and in whole mounts^13,14^; and (4) DNA metabolic labeling for assessing S-phase progression^12^. Application of these methods has revealed cellular responses to injury and regeneration. For example, *P. mytiligera* regeneration is characterized by increased cell proliferation as observed by metabolic DNA labeling^12^, and that cells expressing *BMP1* are detectable in the injured tunic 12 hours following injury^14^.

Despite progress in optimization of molecular methods, techniques for isolation of live cells are missing in *P. mytiligera*. Therefore, isolation of cellular populations that are implicated in regeneration, such as stem cells, phagocytes or other immune cells^15–17^, and analysis of their function are limited. Extraction of live cells involves both isolation of cells from tissues, and cell purification using Fluorescence-Activated Cell Sorting (FACS). Cell isolation requires tissue-specific optimization, based on the tissue characteristics, including tissue composition, availability, and state^18–21^. Optimization of both the mechanical dissociation of the tissue and subsequent enzymatic digestion, if applied, is critical for obtaining true representation of the cellular populations that are found in the tissue^18,19^. Severe mechanical dissociation or harsh enzymatic treatment are often detrimental to the sample integrity and viability of sensitive cell types. By contrast, inefficient tissue disruption could result in poor cellular extraction from tissues^19^. Therefore, it is necessary to empirically determine the optimal conditions for tissue dissociation prior to further processing.

Tissue dissociation generates a mix of cell populations, as well as cellular and non-cellular debris. Purification of live cells requires distinguishing between these components. This is often achieved by using FACS to separate cells based on their morphological properties (e.g., particle size and granularity) and by using functional and genetic fluorescent markers. Cells from marine organisms often have naturally occurring fluorescence (i.e., autofluorescence)^22–24^. Therefore, application of fluorescent reagents for FACS requires prior assessment of the inherent fluorescent properties of the sample. Moreover, biological properties of the sample also influence the choice of fluorescent reagents compatible for cell purification^5^.

Cell extraction from *P. mytiligera* is challenging. Adult animals have opaque tissues and are covered by a thick, tough integumentary tissue known as tunic. The tunic is composed of cellulose, collagens and other extracellular-matrix proteins, vasculature, and free cells ^25,26^. The body wall epidermis is tough and highly pigmented, and internal tissues are fibrotic. Moreover, the tunic is covered by epibionts, such as algae and invertebrates^27^. Therefore, efficient cell purification strategies are required for overcoming these challenges.

Here, we developed a method for dissociating *P. mytiligera* tissues and extracting single cells for further analysis. We analyzed the autofluorescence properties of the dissociated tissues using different flow cytometry approaches, and optimized a strategy for isolating live cell populations based on non-species-specific fluorescent markers. We demonstrated the utility of fluorescent labeling reagents on three flow cytometry platforms, including analyzer, sorter, and imaging flow cytometer. Finally, we demonstrate potential uses of this method, by extracting RNA from FACS-purified cells and profiling their gene expression, and by performing proof-of-concept ex vivo phagocytosis assay. Our work facilitates cell purification from *P. mytiligera*, which is necessary for development of functional cell assays, and therefore advances the analysis of the cellular basis of *P. mytiligera* regeneration.

## Results

### Cell extraction from *P. mytiligera* tissues

To isolate cells from adult *P. mytiligera*, we surgically separated the covering tunic from the underlying tissues. We isolated tissues from the anterior body region, including the neural complex, oral and atrial siphons, branchial basket, and the surrounding body wall (Fig 1A-C). Isolated tissues (<3 cm long) were diced to small fragments (∼1 mm) using a razor on ice (Fig 1D-E). Then, the fine fragments were washed in medium (Methods). The fragments were then subjected to enzymatic treatment and filtered using a mesh. Alternatively, fragments were filtered using a mesh without enzymatic treatment. Cells were collected by centrifugation, and additional filtration steps, resulting in a pigmented cell suspension (Fig 1F-H; Methods). Finally, cells were labeled with viability dye (propidium iodide, PI), and cell viability was assessed by microscopy. We tested different medium conditions for optimizing viability of the extracted cells (Methods). The media tested included artificial seawater (ASW) recipes, which were previously used for marine invertebrate cell isolation^28,29^ or PBS, in a range of pH and media osmolarities. Cell viability in most media conditions was very poor, with >90% non-viable cells, as indicated by positive PI labeling (Methods). Mild enzymatic treatment with collagenase I and no-enzymatic treatment resulted in the best viable cell recovery (Fig 1I-J; Methods). However, omitting enzymatic treatment has reduced the length of the procedure, and it was therefore favored for maintaining cell health^18^.

**Figure 1.**
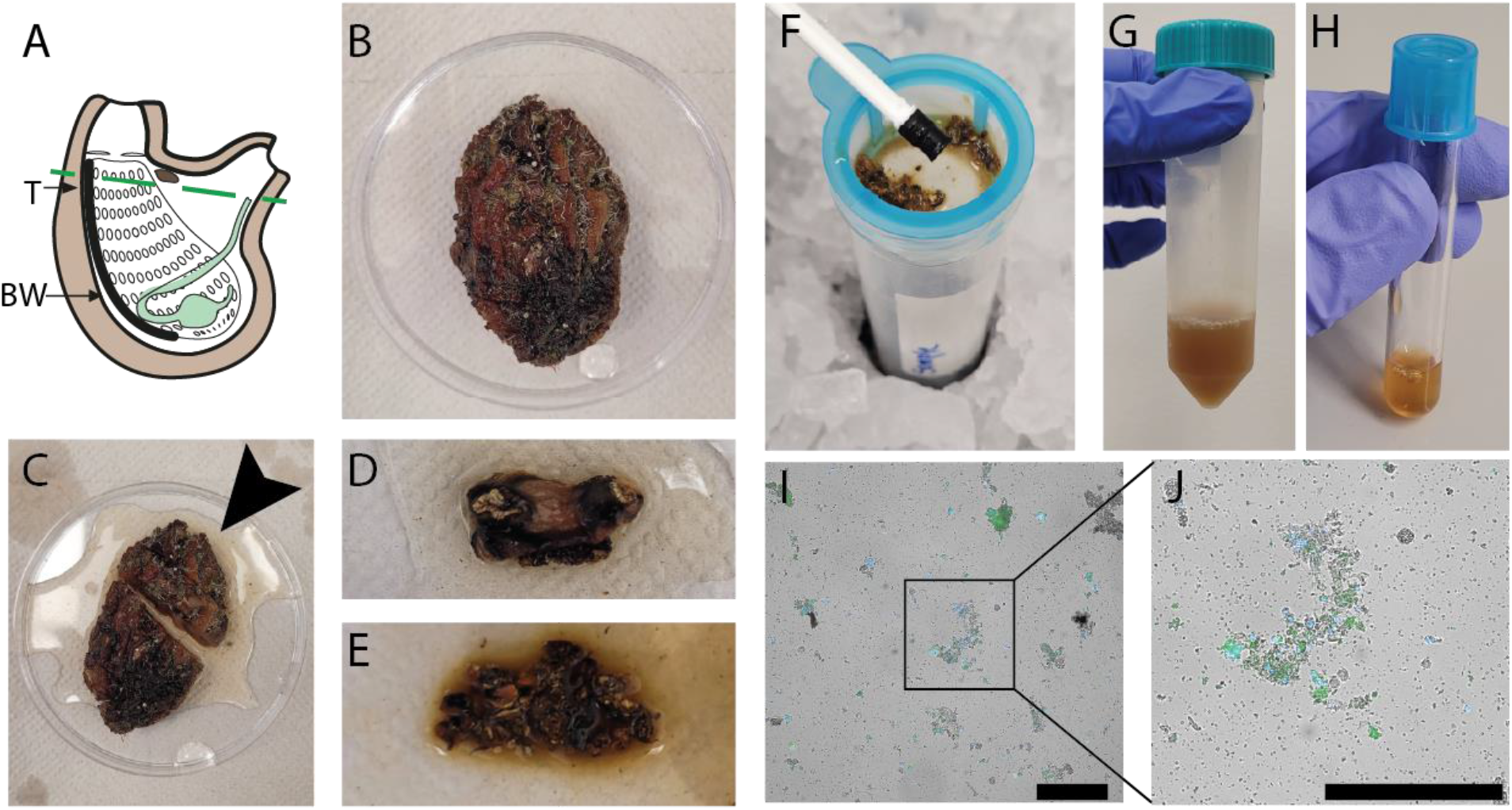
Optimization of cell extraction from *P. mytiligera*. (A) Diagram of adult *P. mytiligera* showing the body region isolated for cell extraction (region above the dashed line). T, tunic; BW, body wall. (B) Photo of adult *P. mytiligera* before dissection showing the opaque tunic and covering epibionts. (C) Separation of anterior region (black arrow) for cell isolation, prior to removal of covering tunic. (D) Isolated anterior region following tunic removal. (E) Tissue fragments following fine dicing using a razor blade. (F) Mechanical filtration of finely diced fragments with a 40 µm filter using a plunger. (G-H) Resultant cell suspension prior to (G) and following (H) repeated centrifugation for cell collection. (I-J) Live cells obtained using optimal cellular extraction parameters. Cells are labeled with nuclear (Hoechst) and viability (calcein) labels, blue and green, respectively. Debris and cell aggregates are detectable as well. Square (I) indicates higher magnification of a region containing cells shown in panel J. Scale = 50 µm.

### Analysis of autofluorescence of *P. mytiligera* cells

Isolated cells from ascidians frequently display autofluorescence^22–24^. To assess the extent of autofluorescence in *P. mytiligera*, we analyzed unlabeled cells extracted from tissues using flow cytometry and microscopy (Methods). Flow cytometry indicated that a small subset (<1%) of the unlabeled cells extracted from the upper body region, without the tunic, has detectable autofluorescence in the tested wavelengths (Fig 2A). This indicated that many fluorophores were likely compatible with *P. mytiligera* cells for cell purification. Similarly, we assessed autofluorescence in extracts from the tunic of *P. mytiligera* (Fig 2B). We found higher levels of autofluorescence in every tested wavelength, and particularly in the far-red range (excitation/emission, 638/660 nm), where 4.21% of the cells displayed autofluorescence (average of three replicates; Fig 2B).

**Figure 2.**
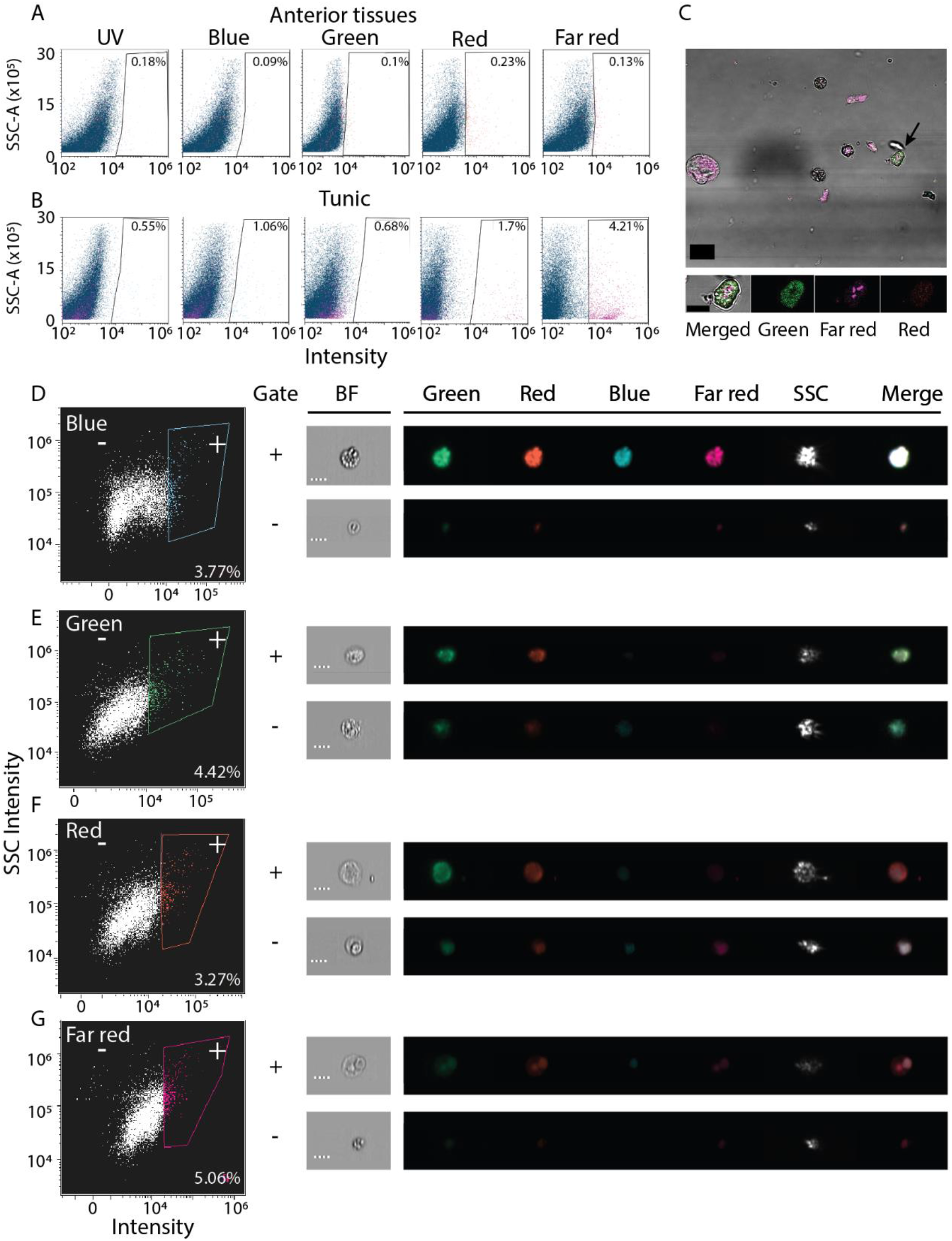
Autofluorescence in isolated *P. mytiligera* cells. (A-B) Shown are representative flow cytometry plots of cells isolated from anterior tissues (A) or tunic (B) selected from three independent replicates of the experiment. X axis shows intensity measured using laser and filter corresponding to the label and Y axis is the measured side scatter. Gates defining cells having autofluorescence are shown with the fraction of cells detected in the gate out of all cells. The label is the average of three experiments. (C) Shown are cells having autofluorescence imaged using fluorescence microscopy (Methods). Tissue was dissociated and imaged without fluorescent labeling. Black arrow (top panel) indicates the cell in the high magnification image (bottom). scale = 20 and 10 µm, top and bottom panels, respectively. (D-G) Autofluorescence analysis using an imaging flow cytometer (Methods). Left panels show representative flow cytometry analyses. Right panels: shown are images of cells registered during the ImageStream analysis. Shown is the fluorescence intensity in every channel and a bright field image (BF). Bounding rectangles were added to the captured images for clarity. Plus and minus symbols represent inclusion or exclusion in the gate, respectively. Label indicates the fraction of cells included in the positive (plus) gate. Scale = 7 µm.

The detected autofluorescence could originate from a distinct cell population. We used two strategies to validate the presence of cells having autofluorescence. First, we imaged unlabeled dissociated cells from the body wall, and detected autofluorescence in both the green and far-red channels (Fig 2C; Methods). These cells appeared granular, in agreement with previous reports of autofluorescence from granular ascidian cells^22,24^. Second, we used a platform combining flow cytometry and imaging (ImageStream; Methods) for detecting autofluorescence in cells in unlabeled body wall samples (Fig 2D). In agreement with traditional flow cytometry and microscopy (Fig 2A, C), we detected autofluorescence in cells, although in this analysis their abundance was higher (Fig 2D-G). This higher prevalence could reflect differences in sensitivity of the instrument, or availability of filters and excitation lasers for each instrument. In this analysis, registered events were imaged during detection by the flow cytometer (Fig 2D-G).

### Optimization of purification of live cells using FACS

Purification of live cells from a dissociated tissue requires elimination of debris and dead cells. We dissociated *P. mytiligera* tissues to cell suspension (Methods). Prior to cell purification we found an abundance of cellular debris and cell aggregates (Fig 1J-I). We separated debris and removed cell aggregates by using side scatter area (SSC-A), and forward scatter area and height, FSC-A and FSC-H, respectively (Fig 3A-B). We used labeling reagents to separate live and dead cells. First, we used calcein to detect live cells (Fig 3C). Then, we labeled nuclei using DRAQ5 and Hoechst for detecting nucleated particles (Fig 3D-G). Comparison of unlabeled (Fig 3D, F) and labeled samples (Fig 3E,G) showed strong enrichment (>10-fold) for nucleated particles following labeling. Combining Hoechst and calcein was particularly useful for isolating live cells (Fig 3H), and distinguished between at least three potential cell populations. We isolated the potential three cell populations using FACS and imaged the cells using confocal microscopy (Methods; Fig 3H-I). We found that the two populations showing high intensity of calcein emission were indeed cells, based on assessment of nuclear and cytoplasmic morphologies and size (Fig 3I). By contrast, cells having low Hoechst and low calcein, were very sparse on the slide. This population did not have a distinct morphology of viable live cells, which suggested having poor health or indicated of cell loss prior to imaging by microscopy (Fig 3I).

**Figure 3.**
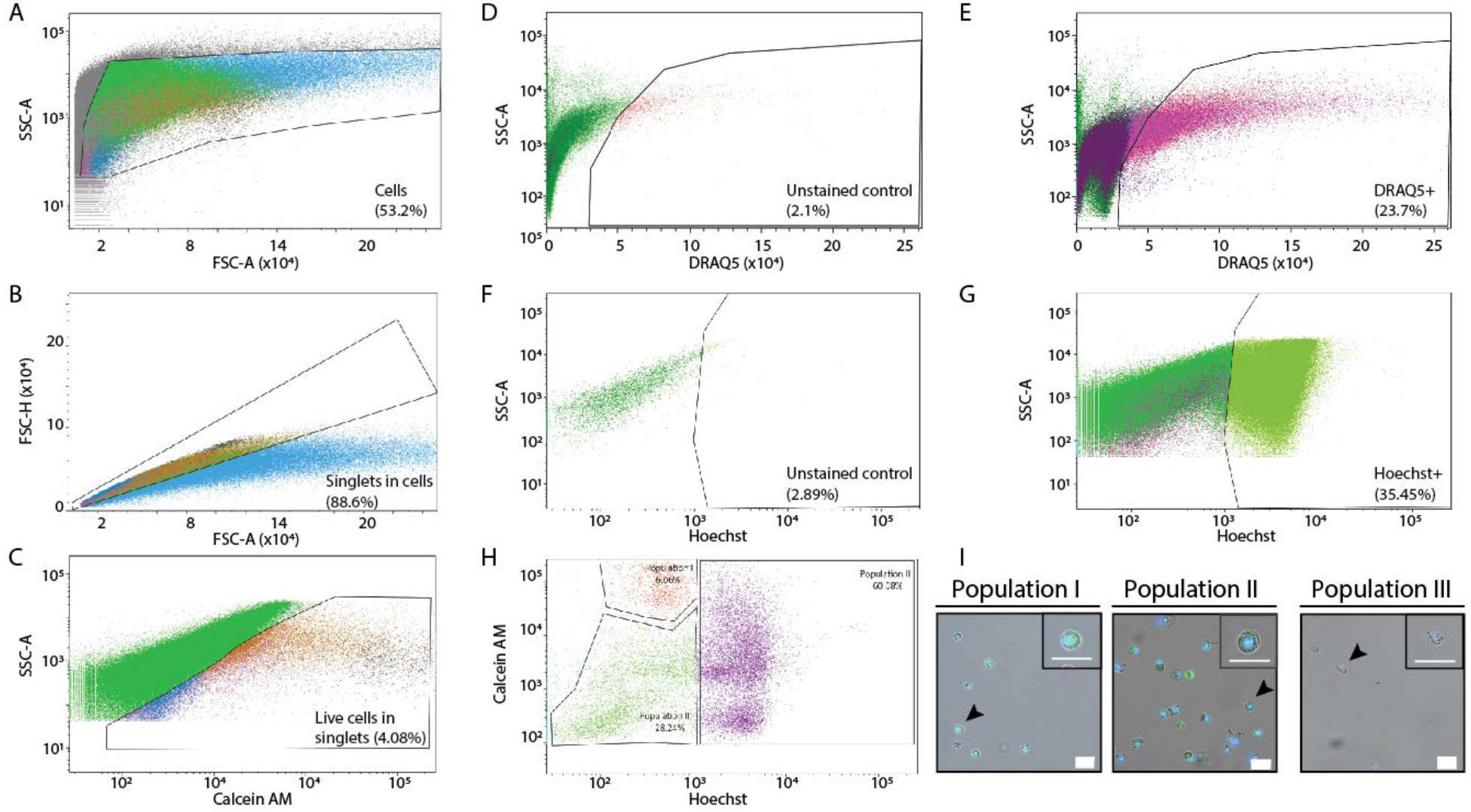
Purification of *P. mytiligera* cells by FACS. (A-H) Shown are flow cytometry analyses in labeled and unlabeled samples. X and Y axis show signal intensity (arbitrary units, A.U.) as measured by FACS in the channel shown in the label (Methods). Filters for different channels and wavelength were as follows: Hoechst: 405 nm, 450/40 band pass (BP), calcein green (AM): 488nm, 502 long pass (LP) 530/30 BP, calcein Violet: 405nm, 450/40 BP, and DR: 640 nm, 690 LP 730/45 BP. (A) Gating strategy for eliminating non-cellular particles using side scatter area (SSC-A) and forward scatter area (FSC-A). (B) Separating aggregates and large particles from single cells (singlets) using FSC-A and FSC height (FSC-H). (C) Using calcein labeling and SSC-A for isolating live cells from the singlet channel. (D-E) Analysis of signal in the DR emission wavelength in unlabeled (D) and DR-labeled (E) samples shows over 10-fold enrichment of live single cells (singlets) in the labeled sample. (F-G) Analysis of signal in the Hoechst emission wavelength in unlabeled (F) and Hoechst-labeled (G) samples shows over 12-fold enrichment of live single cells (singlets) in the labeled sample. (H-I) Detection of putative cell populations by combining Hoechst and calcein labeling. The cells from the different gates (H) were sorted by FACS. Then, the purified samples were examined using confocal microscopy (I; Methods). Samples having high Hoechst and/or calcein (population I and II) showed an abundance of cells with little debris. By contrast, cells having low Hoechst and calcein in the flow cytometry analysis (population III) were sparse and showed poor morphology. Scale = 20 µm.

### Application of imaging flow cytometry for detection of cell types

We applied the labeling strategies described above and used an imaging flow cytometer (Fig 4A; Methods). The instrument captures bright field and fluorescence images for the analyzed flow cytometry events. We manually analyzed the images and extracted photos of cells with distinct morphologies, which resembled known ascidian cell types (Fig 4B)^30–33^. We observed enrichment in cell morphologies that are typical to hematopoietic cell types, suggesting an enrichment for such cell types in this flow cytometry approach. This could be the outcome of a less complex morphology or better compatibility of such cells with the tissue dissociation protocol. We also found cells having morphologies, which we could not associate with known cell types (Fig 4C). This analysis indicated that our approach indeed recovered multiple cell types with an intact cell morphology.

**Figure 4.**
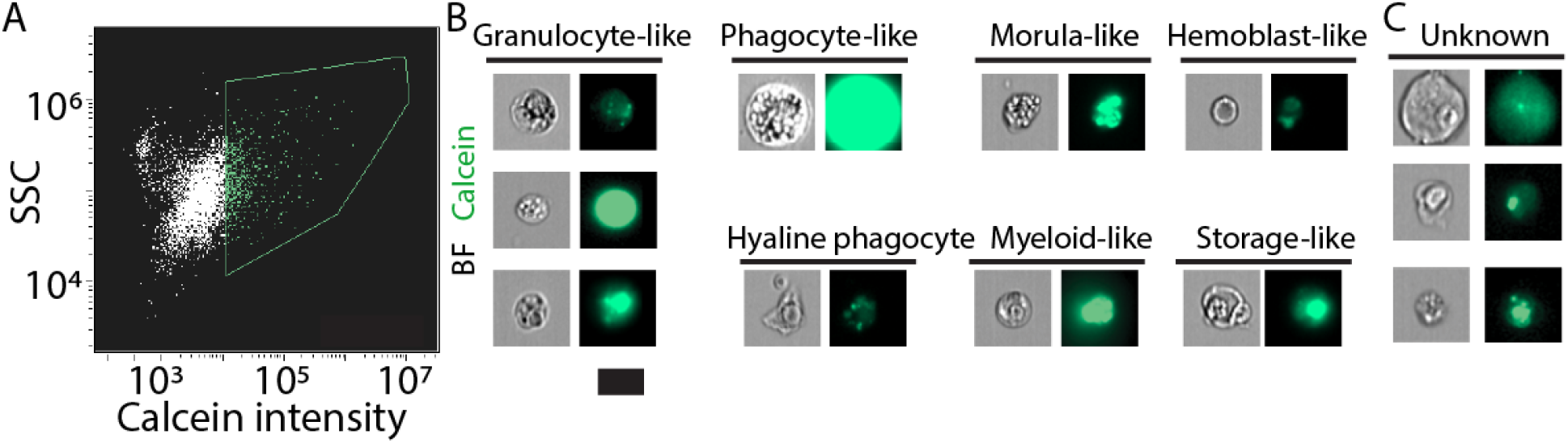
Live cell detection using an imaging flow cytometer. (A) Live cells were detected using calcein labeling and assessment of SSC (Methods). Green contour shows gate used for selecting the calcein positive cells. (B-C) Images of the analyzed cells were acquired by the instrument. (B) Shown are cells that have morphology that resembled known ascidian cell types. Enrichment for hematopoietic cell types was observed, potentially because of preferential isolation of these cells. (C) Shown are cells that we did not classify to a cell type. Scale = 10 µm.

### RNA analysis from sorted *P. mytiligera* cells

We next tested whether the FACS-purified cells are compatible with standard RNA extraction. *P. mytiligera* tissues were dissociated. Then, cell suspensions were subjected to FACS purification directly into the extraction buffer, TRIzol LS (Methods). RNA extraction yielded high quality RNA, even from a small number of cells (<10k; Fig 5A). To test whether the isolated RNA represented gene expression from a diversity of *P. mytiligera* tissues, we prepared RNA sequencing (RNAseq) libraries from isolated RNA. Following sequencing and mapping to the *Polycarpa* transcriptome assembly^13,14^, we tested whether the gene expression represented cells from multiple tissue types (Methods). We assessed the expression levels of gene markers associated with different cell and tissue types (e.g., muscle, neurons, ciliated cells). We found that tissue specific genes are indeed expressed in our RNAseq libraries (Fig 5B). However, the small number of cells used for RNA extraction of each library has likely contributed to variability in the observed gene expression. These results indicate that RNA from FACS-purified cells, using this approach, could be used for analyzing gene expression, and therefore facilitate the characterization of different cell populations.

**Figure 5.**
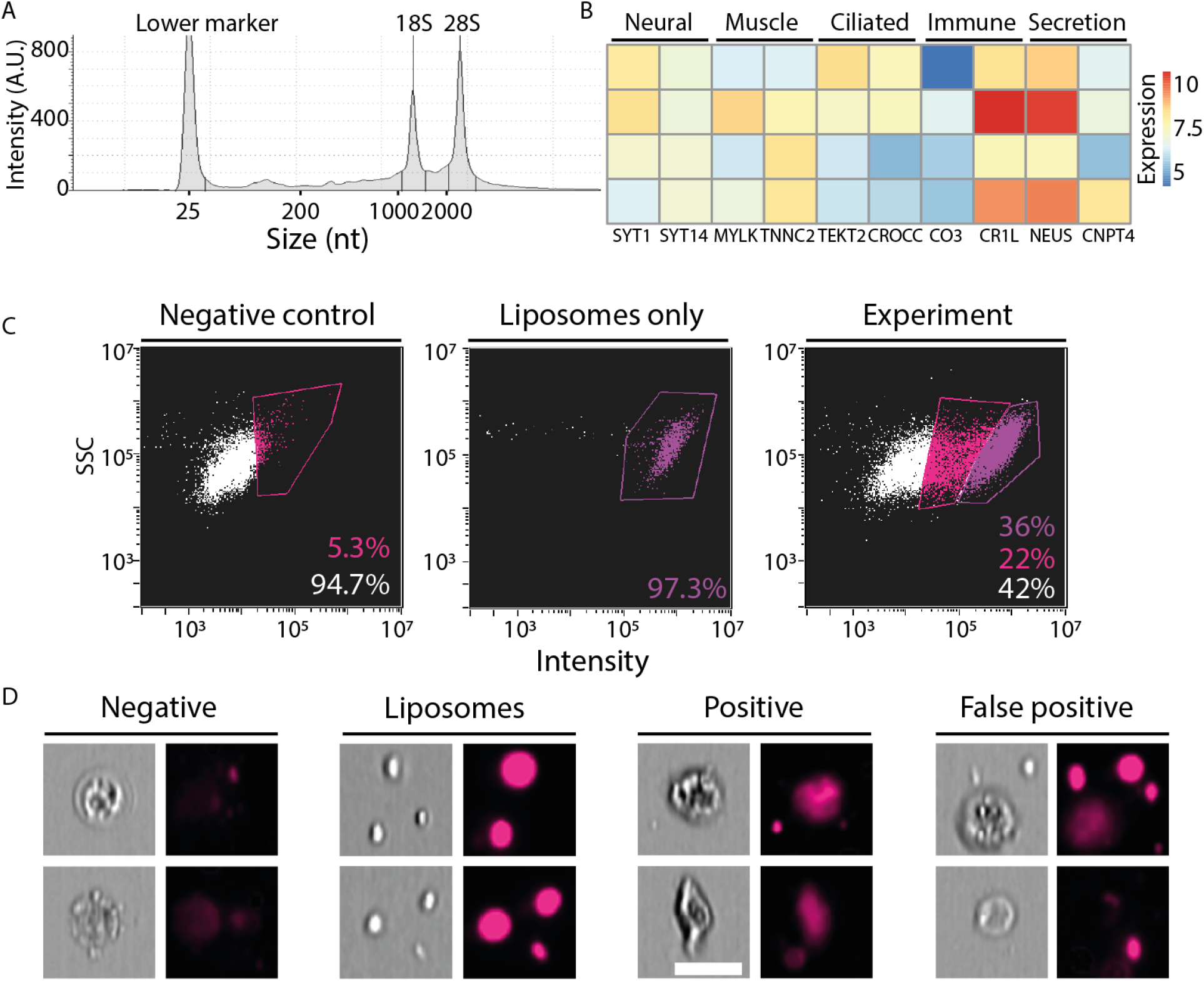
Application of FACS. (A) Shown is a representative microcapillary electrophoresis analysis of RNA that was extracted from ∼10,000 cells FACS-purified cells (Methods). A histogram of RNA species abundance showed that the 18S and 28S ribosomal RNAs remained intact, indicating that the integrity of the isolated RNA from the purified cells was high. (B) Expression of genes that have been associated with expression in different tissues, in other model systems, is shown. Blue and red, low and high gene expression, respectively. Expression was calculated using the variance-stabilizing transformation in DESeq2^34^. Rows represent independent RNAseq libraries that were prepared from purified cells, columns represent genes, and are annotated with gene symbols. (C) Flow cytometry plots showing fluorescence in far red (642-745 nm) and SSC, X and Y axes, respectively, in three samples: left, negative, cells that were not incubated with fluorescent liposomes; middle, liposomes without cells; right, experiment, cells incubated with fluorescent liposomes. The negative (unlabeled) and liposome only samples were used for determining the threshold of true and background signal. (D) Representative images from the imaging flow cytometer showing events captured in this analysis. Negative: cells showing low fluorescence in the far-red channel following incubation with fluorescent liposomes. Liposomes: aggregates of liposomes detected by the imaging flow cytometer. Positive: cells incubated with fluorescent liposomes showing high intensity signal localized to the cell body. False positive: cells detected as positive for high intensity signal, and that by inspection of the photo were found to have the liposomes attached externally to the cell body. Scale = 10 µm.

### In vitro phagocytosis assay using extracted *P. mytiligera* cells

We next tested whether our cell extraction protocol and flow cytometry approach could be used for functional cell assays. We performed a proof-of-concept ex vivo phagocytosis assay. We incubated extracted *P. mytiligera* cells with fluorescent liposomes, containing DiD fluorophore (far red; excitation/emission: 644/665 nm), which could be ingested only by phagocytosis (Methods). Following incubation, cells were analyzed using an imaging flow cytometer (ImageStream; Methods). Background fluorescence level was determined using an unlabeled sample (Fig 5C-D), and fluorescence of free liposomes was determined by using a cell-free sample containing only media and fluorescent liposomes. The imaging flow cytometer acquired photos of: (1) negative cells: cells negative for far red fluorescence; (2) positive cells: cells having far red fluorescence localized in the cell body; (3) false positives: cells determined as positive by the flow cytometer, according to fluorescence intensity. However, manual inspection of the photo indicated that the liposomes were external to the cell body; (4) free floating liposomes. Further optimization of this assay could reduce the amount of free and cell-attached liposomes. Our results suggested that a fraction of the cells from the anterior body region could uptake large liposomes, and therefore might represent a phagocyte population.

## Discussion

In this study, we developed a method for dissociating *P. mytiligera* tissues and purifying live cells using FACS for further processing. We used this approach for analyzing cells on different flow cytometry instruments, including an analyzer, a sorter, and an imaging flow cytometer. These instruments represent widespread approaches for cell analysis and purification by flow cytometry.

Optimization of tissue dissociation and cell purification required overcoming challenges inherent to this species anatomy, but also relevant to many ascidians species and other marine invertebrates. The thick tunic restricts internal tissue isolation, and body wall tissues are difficult to dissociate without damaging cells. We therefore tested different media conditions for tissue dissociation and examined the disassociation outcome by microscopy. Cellular extraction using enzymatic dissociation of tissue fragments was inconsistent when using high enzyme concentration. This might be a result of variability in tissue composition, as the primary tissues were isolated from different specimens. Alternatively, that might be a result of harsh enzyme activity that affected cell viability. The use of manual mechanical cell extraction resulted in better consistency in cell extraction in our hands, and we therefore used it in subsequent experiments.

Development of a FACS approach for cell purification required autofluorescence analysis, which has been documented in various ascidians^22–24^. We found that commonly used reagents were applicable to *P. mytiligera*, following task-specific optimization. Combination of Hoechst with calcein was particularly effective for isolating live cells, and detection of multiple putative cellular populations. This approach could be applied in future studies to assess the identity and function of uncharacterized cell populations. Further optimization of Hoechst labeling could be utilized in the future to separate cells based on their DNA content^35^, as a proxy for isolating cells based on their cell cycle state. This would be extremely useful for understanding the cellular basis of regeneration^12^. In planarians, highly regenerative flatworms, pluripotent stem cells mediate regeneration^36,37^. In *P. mytiligera*, regeneration involves cell proliferation^12^, but the cells that promote this process have not been identified yet. Therefore, the methods developed here facilitate studying the mechanisms of regeneration in the *P. mytiligera* system.

We showed here two proof-of-concept experiments that require tissue dissociation and cell purification. First, we applied RNAseq to FACS-purified cells. Genes likely expressed in different cell types were represented in the dataset. However, the libraries were prepared from RNA extracted from a small number of cells (∼10k), and therefore were likely not to capture the true complexity of the analyzed tissues. Preparation of RNAseq libraries using larger amounts of input RNA, or the use of alternative approaches (e.g., single cell RNAseq), should be considered for comparative gene expression analyses of *Polycarpa* purified cells. In a second proof-of-concept experiment, we analyzed the cellular uptake, ex vivo, of fluorescent particles. Comparison of cells incubated with the reagent with control cells showed a major increase in signal intensity in the treated cells. This suggests that a similar assay could be used for studying phagocytosis in *P. mytiligera*.

The approach and method we developed here could be further tested in other ascidians, and represents a significant step towards establishment of cell function assays in this emerging model system.

## Methods

### Animal collection and maintenance

Animals were collected by SCUBA diving in the bay of Aqaba (Eilat) and transferred to Tel Aviv University. Animals were maintained in a recirculating aquarium system with artificial sea water (Red Sea Salt, 8 kh) mixed in reverse osmosis water.

### Media used for testing tissue dissociation

The following media compositions and enzymes were tested for *P. mytiligera* tissues dissociation: (1) NaCl (449mM), Na2SO4 (33mM), KCl (9mM), NaHCO3 (2.15 mM), EDTA (292.24 mM), Tris-Cl (5ml), pH range tested between 8-8.2, concentration part per trillion (ppt) 32. This media was used in combination with trypsin (0.25%, Sartorius, 03-052-1A) and collagenase (1 mg/ml, Sigma, C0130) added 1:10. (2) NaCl (0.4M), KCl (10mM), MgSO4-7H2O (7.6mM), MgCl2-6H2O (52mM), Na2SO4 (21mM), NaHCO3 (3mM), SrCl2 (0.17mM), EDTA (5mM). pH 8, ppt 40. This media was tested by adding collagenase (1 mg/ml) to 1:10 and 1:20. (3) PBSx3.3, pH 7.5, ppt 35. This media was tested by adding collagenase (1 mg/ml) 1:20 and trypsin (0.25%). The efficiency of the different combination of media and enzymatic activity was determined by analyzing the percentage of viability dye (propidium iodide, PI) (1:1000) positive cells using hemocytometer.

### Optimized tissue dissociation procedure

Tissue fragments of up to three centimeters were isolated surgically from adult animals using a blade. Following tunic removal, the tissue fragments were washed with filtered (0.2 μm) artificial sea water (ASW; 0.4M NaCl, 10mM KCl, 7.6mM MgSO4-7H2O, 52mM MgCl2-6H2O,21mM Na2SO4, 3mM NaHCO3, 0.17mM SrCl2, 10 mM HEPES, 5mM EDTA, pH 8, 40 ppt) and transferred into a sterile petri dish on ice. Using a blade, the tissue fragments were cut to smaller fragments, centrifuged at 300G at 4 °C for 5 minutes and resuspended in 1 ִml artificial sea water.

For the enzymatic treatment samples were incubated with enzymes (collagenase I, Sigma, C0130) for 7 min at room temperature (RT) and mixed by pipetting. Then 1 ml of cold ASW with 0.5% BSA (Mercury, 821006) was added to the sample. The sample was then filtered through a 40 μm mesh filter (LifeGene, G-CSS010040S). Cell suspensions were further dissociated on the mesh by using a sterile plunger of 1 ml syringe (PIC, 00603308), washed, and collected in ASW. Then, cells were collected by centrifugation at 300G at 4 °C for 7 minutes, and then resuspended in 1 ml ASW with 0.5% BSA. Cells were labeled using the nuclear dyes, Hoechst 33342 (Thermo Fisher, H3570) and DRAQ5 (Abcam, ab108410), to a final concentration of 20 μM.

Calcein-AM (BioLegend, #425201) and Calcein Violet-AM (BioLegend, # 425203) were used as live cell labeling dye by supplementing to a final concentration of 4 μM. Cells were incubated for 30 minutes at room temperature. Cell concentration was estimated using a hemocytometer on an inverted confocal microscope system (Zeiss LSM800).

### Cell sorting, Flow cytometry analysis and live cells microscopy

FACS reading was done on BD FACS Aria II, and gating for cell sorting was done using the BD FACSDiva™ software (Becton, Dickinson Biosciences). Flow cytometry analysis was done using Beckman Coulter CytoFLEX 4L(Beckman Coulter). Imaging on flow cytometry was performed using the ImageStream^x^ mk II platform (Luminex). For determining the gating of cells and debris, sorting and observation by confocal microscopy was done several times. Analysis of flow cytometry data was done using Kaluza Analysis Software (Beckman Coulter) and IDEAS® 6.2 ImageStream Analysis Software. Specification of excitation laser and optical filter for emission for each analysis is shown in the figures. The excitation laser was measured in nm and filters are stated as long pass (LP) and band pass (BP).

Cell sorting was performed with 100 μm nozzle size and sorted directly into Eppendorf tubes containing 100 μl of ASW in order to minimize cellular stress. Cells (10,000–50,000) of each population of interest were sorted at a speed of 1500 cells / second.

For live cell microscopy, sorted cells were collected by centrifugation at 4°C; 300G; 10 min, the cells were resuspended in 20 μl of ASW and counted using a hemocytometer. Images of sorted cells were acquired using confocal microscopy (Zeiss LSM800) using the Zeiss Zen Blue v2.3 software. In addition, fluorescence microscopy was used to image autofluorescence in unlabeled cells (Leica SP8, Laser: 488 Filter: 500-560; Laser: 561 Filter: 571-639; Laser: 633 Filter: 645-730).

### Phagocytosis assay

Cells were extracted from the anterior body region of animals following tunic removal. Extracted cells were resuspended in 800 µl of L-15 media containing glucose and Fetal Bovine Serum (Sigma Aldrich, F4135) in a 24-well plate. For the treatment groups, the media was supplemented with 8 μL of liposomes containing the fluorophore DiD (Encapsula Nanosciences, CLD-8904), with excitation and emission spectra 644 and 665 nm, respectively. Incubation in media containing fluorescent liposomes was performed for 4.5 hours at room temperature in the dark. Prior to analysis using an imaging flow cytometer (ImageStream MKII), the media was supplemented with 4′,6-diamidino-2-phenylindole (DAPI) for 30 minutes (1 mg/µl).

### RNA quality assessment and extraction

RNA extractions were performed using TRIzol LS (Thermo Fisher; #10296010), following the manufacturer’s protocol. RNA concentration was measured by RNA fluorometry (Qubit 4, Life Technologies) using the RNA high sensitivity kit (Q32852). RNA integrity was assessed using a microcapillary electrophoresis (TapeStation 4200, G2991BA) with RNA ScreenTape kit (Agilent Technologies, #5067-5576).

### RNA sequencing library preparation

Illumina-sequencing compatible RNAseq libraries were prepared using New England Biosciences (NEB) NEBNext Ultra II Directional RNA mRNA seq library (NEB; #E7760L) according to the manufacturer protocol, with at least 10 ng total RNA per library, and an estimated number of cells in the range of 10,000. Briefly, mRNA was enriched by polyA selection using poly-dT paramagnetic beads included in the kit. Then, RNA was fragmented according to the protocol, and complementary DNA (cDNA) was produced by reverse transcription. Following second-strand synthesis, the resultant DNA was end-repaired. Adapter ligation was performed according to protocol. The libraries were then amplified by PCR using barcoded primers according to the kit instructions. Quantitative PCR (qPCR) with Illumina primers was used to determine the optimal number of PCR amplification cycles. The number of amplification cycles (17-20) represented an early exponential increase in the qPCR signal. Following PCR amplification, libraries were sequenced using Novogene sequencing services.

### RNAseq libraries analysis

RNAseq libraries were processed similarly to previously described^14^. Briefly, RNAseq libraries were trimmed using cutadapt v2.85 using the Illumina adapter sequences^38^. Then, the preprocessed library reads #1 (first read in pair) were mapped to a transcriptome assembly of *P. mytiligera* using bowtie2 v2.4 using flags -local --trim3 80 --trim5 2 --sensitive-local^39^. A gene expression matrix was produced using featureCounts package v2.0.0 with parameters allowing multi mapping [-M -s 0]^40^. Gene expression levels in the libraries were estimated using DESeq2 and expression levels, across libraries, were normalized using the variable stabilizing transformation in the DESeq2 package. Genes expressed in a cell type-enriched or tissue-enriched manner were extracted from the GeneCards database manually using tissue and cell type keywords^41^, and similar sequences in *P. mytiligera* were determined by using BLAST search for the best blast hit with e-value < 10^−20^ ^42^. RNAseq data was deposited in the Sequence Read Archive (SRA)^43^ under accession PRJNA996490.

## Author Contributions

T.G., N.H. and O.W. designed the project. T.G. and N.H. performed the experiments. T.G. optimized the tissue dissociation method and applied flow cytometry analysis and sorting. N.H. performed the imaging flow cytometry experiments. N.H. performed the proof-of-concept phagocytosis assays. T.G. prepared RNAseq libraries from FACS-purified cells. O.W. analyzed the RNAseq data. All authors wrote and edited the manuscript.

## Acknowledgements

We thank Dr. Benyamin Rosental from Ben Gurion University for discussions and advice on cell dissociation and flow cytometry method optimization. We thank the Inter-University Institute (IUI) for their support and facilities. We acknowledge Dr. Irena Shur, Dr. Daria Makarovsky, Dr. Sasha Lichtenstein, and Dr. Rami Khosravi from the interdepartmental services at Tel Aviv University’s Faculty of Medicine for assistance with flow cytometry operation and analysis. We thank Dr. Hila Kobo for assistance with Illumina sequencing library preparation. We thank the Wurtzel lab members for comments and discussions. O.W. is supported by the Israel Science Foundation (grant 2039/18) and the European Research Council (no. 853640). O.W. is a Zuckerman Faculty Scholar. T.G. was supported by an excellence postdoctoral scholarship from the Israeli Ministry of Science and by an excellence scholarship from the Tel Aviv University Rector’s office.

